## Supplementary material for "A Fast Approach for Structural and Evolutionary Analysis Based on Energetic Profile Protein Comparison": Supplementary Information.pdf

### Supplementary Tables

**Supplementary Table 1.** The accuracy and computation time for 1-NN classifier based on GR-Align, RMSD, TM-score, Yau-Hausdorff distance, TM-Vec, and the distance between profiles of energy SPE and CPE as a measure of protein dissimilarity.

| **Method** | **Accuracy** | **Time** |
| --- | --- | --- |
| **GR-Align** | 62.3% | 2 min |
| **RMSD** | 59.2% | 1 h |
| **TM-Score** | 61.5% | 9 h 20 min |
| **YH (10 Rotation)** | 70.8% | 10 min |
| **YH (2500 Rotation)** | 81.5% | 4h 10 min |
| **TM-Vec** | 100% | 67 sec |
| **CPE** | 100% | 1 sec |
| **SPE** | 99% | 187 sec |

**Supplementary Table 2.** Comparison of clustering results using Adjusted Rand Index (ARI).

| Method | Type | Cut Tree | ARI |
|---|---|---|---|
| **CPE** | Sequence-Based | 4 | 0.95 |
| **MSA** | Sequence-Based | 3 | 0.49 |
| **TM-Vec** | Sequence-Based | 5 | 0.87 |
| **SPE** | Structure-Based | 3 | 1.00 |
| **RMSD** | Structure-Based | 6 | 0.73 |
| **TM-Score** | Structure-Based | 4 | 0.56 |

**Supplementary Table 3.** Comparison of Clustering Methods by Sequence and Structure: Time, ARI, and Classification Error Across Clusters.

| Based | Method | Time | ARI | | | | Class Error | | | |
|---|---|---|---|---|---|---|---|---|---|---|
| | | | 3 | 4 | 5 | 6 | 3 | 4 | 5 | 6 |
| Sequence | CPE | **0.9 sec** | 0.50 | **0.95** | 0.94 | 0.92 | 0.22 | **0.08** | 0.1 | 0.11 |
| | TM Vec | 89 sec | 0.16 | 0.48 | 0.87 | **0.86** | 0.34 | 0.22 | 0.12 | **0.14** |
| | MSA (ClustalW) | 72 sec | **0.49** | 0.49 | 0.32 | 0.32 | **0.22** | 0.22 | 0.29 | 0.29 |
| Structure | SPE | **3 min** | **1** | **0.95** | 0.93 | 0.66 | **0** | **0.08** | 0.11 | 0.26 |
| | RMSD | 70 min | 0.50 | 0.50 | 0.36 | **0.73** | 0.22 | 0.22 | 0.22 | **0.17** |
| | TM score | 9.7 h | 0.50 | **0.56** | 0.40 | 0.40 | 0.22 | **0.17** | 0.24 | 0.24 |

**Supplementary Table 4.** The PDB IDs of spike proteins.

| pdbID | length | virus | pdbID | length | virus | pdbID | length | virus | pdbID | length | virus |
|---|---|---|---|---|---|---|---|---|---|---|---|
| 6XM5A | 1057 | SARS-CoV-2 | 6CRZA | 1068 | SARS-CoV | 6Q04B | 1159 | MERS-CoV | 6ZGFC | 1060 | SARS-CoV-2 |
| 6ZP2A | 1097 | SARS-CoV-2 | 6CRWA | 1068 | SARS-CoV | 6ACKB | 1065 | SARS-CoV | 6ZGEC | 1098 | SARS-CoV-2 |
| 6ZP1A | 1017 | SARS-CoV-2 | 5W9JA | 463 | MERS-CoV | 6ACJB | 1065 | SARS-CoV | 6Z97C | 991 | SARS-CoV-2 |
| 6ZP0A | 1030 | SARS-CoV-2 | 5XLRA | 1022 | SARS-CoV | 6ACGB | 1065 | SARS-CoV | 7BYRC | 973 | SARS-CoV-2 |
| 6ZOZA | 1069 | SARS-CoV-2 | 5X5FA | 1141 | MERS-CoV | 6ACDB | 1065 | SARS-CoV | 6X6PC | 1017 | SARS-CoV-2 |
| 6ZOYA | 1021 | SARS-CoV-2 | 5X58A | 1054 | SARS-CoV | 6ACCB | 1065 | SARS-CoV | 6Z43C | 991 | SARS-CoV-2 |
| 6ZOXA | 1017 | SARS-CoV-2 | 5WRGA | 736 | SARS-CoV | 5W9OD | 463 | MERS-CoV | 6X2CC | 971 | SARS-CoV-2 |
| 6XEYA | 1034 | SARS-CoV-2 | 6ZP7B | 930 | SARS-CoV-2 | 5W9ND | 463 | MERS-CoV | 6X2AC | 961 | SARS-CoV-2 |
| 6ZGIA | 1098 | SARS-CoV-2 | 6ZP5B | 943 | SARS-CoV-2 | 5W9MD | 457 | MERS-CoV | 6X29C | 972 | SARS-CoV-2 |
| 6ZGGA | 1069 | SARS-CoV-2 | 6ZOWB | 930 | SARS-CoV-2 | 5W9LB | 726 | MERS-CoV | 6WPTC | 952 | SARS-CoV-2 |
| 6ZGFA | 1060 | SARS-CoV-2 | 6ZHDB | 992 | SARS-CoV-2 | 5W9KD | 462 | MERS-CoV | 6WPSE | 955 | SARS-CoV-2 |
| 6ZGEA | 1098 | SARS-CoV-2 | 6XM5B | 1056 | SARS-CoV-2 | 5W9JD | 463 | MERS-CoV | 6VYBC | 960 | SARS-CoV-2 |
| 6Z97A | 995 | SARS-CoV-2 | 6ZP2B | 1097 | SARS-CoV-2 | 5W9HD | 463 | MERS-CoV | 6VXXC | 972 | SARS-CoV-2 |
| 6XCMA | 966 | SARS-CoV-2 | 6ZP1B | 1017 | SARS-CoV-2 | 5XLRB | 1022 | SARS-CoV | 6VSBC | 973 | SARS-CoV-2 |
| 6X6PA | 1017 | SARS-CoV-2 | 6ZP0B | 1030 | SARS-CoV-2 | 5X5FB | 1141 | MERS-CoV | 6Q07C | 1159 | MERS-CoV |
| 6X2CA | 971 | SARS-CoV-2 | 6ZOZB | 1069 | SARS-CoV-2 | 5X5CB | 1141 | MERS-CoV | 6Q06C | 1159 | MERS-CoV |
| 6X2BA | 963 | SARS-CoV-2 | 6ZOYB | 1021 | SARS-CoV-2 | 5X5BB | 1053 | SARS-CoV | 6Q05C | 1159 | MERS-CoV |
| 6X2AA | 966 | SARS-CoV-2 | 6ZOXB | 1017 | SARS-CoV-2 | 5X58B | 1053 | SARS-CoV | 6Q04C | 1159 | MERS-CoV |
| 6X29A | 972 | SARS-CoV-2 | 6XEYB | 1034 | SARS-CoV-2 | 5WRGB | 736 | SARS-CoV | 6NB3C | 1169 | MERS-CoV |
| 6WPTA | 945 | SARS-CoV-2 | 6XKLB | 976 | SARS-CoV-2 | 6ZP7C | 943 | SARS-CoV-2 | 6ACCC | 1065 | SARS-CoV |
| 6WPSA | 955 | SARS-CoV-2 | 7C2LB | 1055 | SARS-CoV-2 | 6ZP5C | 930 | SARS-CoV-2 | 6CS1C | 1068 | SARS-CoV |
| 6VYBA | 966 | SARS-CoV-2 | 6ZGIB | 1098 | SARS-CoV-2 | 6ZOWC | 943 | SARS-CoV-2 | 6CS0C | 1069 | SARS-CoV |
| 6VXXA | 972 | SARS-CoV-2 | 6ZGHB | 1077 | SARS-CoV-2 | 6ZHDC | 990 | SARS-CoV-2 | 6CRZC | 1069 | SARS-CoV |
| 6Q07A | 1159 | MERS-CoV | 6ZGFB | 1060 | SARS-CoV-2 | 6XM5C | 1036 | SARS-CoV-2 | 6CRXC | 1069 | SARS-CoV |
| 6Q06A | 1159 | MERS-CoV | 6ZGEB | 1098 | SARS-CoV-2 | 6ZP2C | 1097 | SARS-CoV-2 | 6CRWC | 1068 | SARS-CoV |
| 6Q05A | 1159 | MERS-CoV | 7BYRB | 998 | SARS-CoV-2 | 6ZP1C | 1017 | SARS-CoV-2 | 5W9OG | 463 | MERS-CoV |
| 6Q04A | 1159 | MERS-CoV | 6X6PB | 1017 | SARS-CoV-2 | 6ZP0C | 1030 | SARS-CoV-2 | 5W9NG | 457 | MERS-CoV |
| 6NB6A | 1052 | SARS-CoV | 6Z43B | 992 | SARS-CoV-2 | 6ZOZC | 1070 | SARS-CoV-2 | 5W9ME | 726 | MERS-CoV |
| 6NB4A | 1169 | MERS-CoV | 6X2CB | 971 | SARS-CoV-2 | 6ZOYC | 1021 | SARS-CoV-2 | 5W9LC | 726 | MERS-CoV |
| 6NB3A | 1169 | MERS-CoV | 6X29B | 972 | SARS-CoV-2 | 6ZOXC | 1017 | SARS-CoV-2 | 5W9JG | 463 | MERS-CoV |

| | | | | | | | | | | | |
|---|---|---|---|---|---|---|---|---|---|---|---|
| 6ACKA | 1065 | SARS-CoV | 6WPSB | 955 | SARS-CoV-2 | 6XEYC | 1030 | SARS-CoV-2 | 5W9HG | 463 | MERS-CoV |
| 6ACJA | 1065 | SARS-CoV | 6VXXB | 972 | SARS-CoV-2 | 6XKLC | 976 | SARS-CoV-2 | 5XLRC | 1022 | SARS-CoV |
| 6ACGA | 1065 | SARS-CoV | 6VSBB | 973 | SARS-CoV-2 | 7C2LC | 1049 | SARS-CoV-2 | 5X5BC | 1053 | SARS-CoV |
| 6ACDA | 1065 | SARS-CoV | 6Q07B | 1159 | MERS-CoV | 6ZGIC | 1098 | SARS-CoV-2 | 5X58C | 1052 | SARS-CoV |
| 6ACCA | 1065 | SARS-CoV | 6Q06B | 1159 | MERS-CoV | 6ZGHC | 1080 | SARS-CoV-2 | 5WRGC | 736 | SARS-CoV |
| 6CS0A | 1068 | SARS-CoV | 6Q05B | 1159 | MERS-CoV | 6ZGGC | 1067 | SARS-CoV-2 | | | |

**Supplementary Table 5.** The results of 1-NN classification on SARS Proteome using CPE.

| SARS Proteom | Accuracy | Precision | Recall | F1 |
|---|---|---|---|---|
| E_protein | 99,9546 | 100 | 96,55 | 98 |
| N_C-terminaldomain | 100 | 100 | 100 | 100 |
| N_N-terminaldomain | 99,9773 | 99,82 | 100 | 100 |
| NSP1_protein | 99,9773 | 100 | 95,83 | 98 |
| NSP10_protein | 100 | 100 | 100 | 100 |
| NSP12_protein | 100 | 100 | 100 | 100 |
| NSP13_protein | 100 | 100 | 100 | 100 |
| NSP14_protein | 100 | 100 | 100 | 100 |
| NSP15_protein | 100 | 100 | 100 | 100 |
| NSP16_protein | 99,9773 | 99,61 | 100 | 100 |
| NSP2_protein | 100 | 100 | 100 | 100 |
| NSP3_cd21525_SUD_C_SARS-CoV_Nsp3 | 99,9773 | 99,2 | 100 | 100 |
| NSP3_cd21557_Macro_X_Nsp3-like | 99,9546 | 99,19 | 99,19 | 99 |
| NSP3_cd21717_TM_Y_SARS-CoV-like_Nsp3_C | 99,9773 | 100 | 99,19 | 100 |
| NSP3_cd21732_betaCoV_PLPro | 99,9773 | 99,18 | 100 | 100 |
| NSP3_cd21822_SARS-CoV-like_Nsp3_NAB | 99,9319 | 98,33 | 99,16 | 99 |
| NSP3_cl00019_Macro_SF | 99,9773 | 100 | 97,62 | 99 |
| NSP3_cl13138_SUD-M | 99,8638 | 99,17 | 95,97 | 98 |
| NSP3_cl13772_DUF3655 | 100 | 100 | 100 | 100 |
| NSP5_protein | 99,9773 | 100 | 99,59 | 100 |
| NSP7_protein | 100 | 100 | 100 | 100 |
| NSP8_protein | 99,9773 | 99,5 | 100 | 100 |
| NSP9_protein | 99,9092 | 98,21 | 99,4 | 99 |
| orf3a_protein | 100 | 100 | 100 | 100 |
| orf7a_protein | 100 | 100 | 100 | 100 |
| orf8_protein | 100 | 100 | 100 | 100 |
| orf9b_protein | 99,9546 | 91,3 | 100 | 95 |
| S_protein | 100 | 100 | 100 | 100 |
| **Average** | 99,9773 | 99,4111 | 99,375 | 99,4643 |

**Supplementary Table 6.** The results of 1-NN classification on SARS Proteome using SPE.

| SARS Proteom | Accuracy | Precision | Recall | F1 |
|---|---|---|---|---|
| E_protein | 99,9546 | 100 | 96,55 | 98 |
| N_C-terminaldomain | 99,9773 | 100 | 99,82 | 100 |
| N_N-terminaldomain | 100 | 100 | 100 | 100 |
| NSP1_protein | 99,9773 | 100 | 95,83 | 98 |
| NSP10_protein | 99,9773 | 99,55 | 100 | 100 |
| NSP12_protein | 100 | 100 | 100 | 100 |
| NSP13_protein | 100 | 100 | 100 | 100 |
| NSP14_protein | 100 | 100 | 100 | 100 |
| NSP15_protein | 100 | 100 | 100 | 100 |
| NSP16_protein | 100 | 100 | 100 | 100 |
| NSP2_protein | 100 | 100 | 100 | 100 |
| NSP3_cd21525_SUD_C_SARS-CoV_Nsp3 | 99,9092 | 96,88 | 100 | 98 |
| NSP3_cd21557_Macro_X_Nsp3-like | 100 | 100 | 100 | 100 |
| NSP3_cd21717_TM_Y_SARS-CoV-like_Nsp3_C | 100 | 100 | 100 | 100 |
| NSP3_cd21732_betaCoV_PLPro | 100 | 100 | 100 | 100 |
| NSP3_cd21822_SARS-CoV-like_Nsp3_NAB | 99,9546 | 98,35 | 100 | 99 |
| NSP3_cl00019_Macro_SF | 99,7957 | 97,14 | 80,95 | 88 |
| NSP3_cl13138_SUD-M | 99,9773 | 100 | 99,19 | 100 |
| NSP3_cl13772_DUF3655 | 100 | 100 | 100 | 100 |
| NSP5_protein | 100 | 100 | 100 | 100 |
| NSP7_protein | 100 | 100 | 100 | 100 |
| NSP8_protein | 100 | 100 | 100 | 100 |
| NSP9_protein | 99,9546 | 98,81 | 100 | 99 |
| orf3a_protein | 100 | 100 | 100 | 100 |
| orf7a_protein | 99,9546 | 92,86 | 100 | 96 |
| orf8_protein | 99,9773 | 95 | 100 | 97 |
| orf9b_protein | 100 | 100 | 100 | 100 |
| S_protein | 100 | 100 | 100 | 100 |
| **Average** | 99,9789 | 99,2354 | 99,0121 | 99,0357 |

**Supplementary Table 7.** The results of 1-NN classification on SARS Proteome using TM-Vec.

| SARS Proteom | Accuracy | Precision | Recall | F1 |
|---|---|---|---|---|
| E_protein | 99,9773 | 98,31 | 100 | 99 |
| N_C-terminaldomain | 99,9773 | 100 | 99,82 | 100 |
| N_N-terminaldomain | 100 | 100 | 100 | 100 |
| NSP1_protein | 99,9773 | 100 | 95,83 | 98 |
| NSP10_protein | 99,9773 | 100 | 99,55 | 100 |
| NSP12_protein | 100 | 100 | 100 | 100 |
| NSP13_protein | 100 | 100 | 100 | 100 |
| NSP14_protein | 100 | 100 | 100 | 100 |
| NSP15_protein | 100 | 100 | 100 | 100 |
| NSP16_protein | 100 | 100 | 100 | 100 |
| NSP2_protein | 100 | 100 | 100 | 100 |
| NSP3_cd21525_SUD_C_SARS-CoV_Nsp3 | 99,8184 | 94,62 | 99,19 | 97 |
| NSP3_cd21557_Macro_X_Nsp3-like | 100 | 100 | 100 | 100 |
| NSP3_cd21717_TM_Y_SARS-CoV-like_Nsp3_C | 99,9773 | 100 | 99,19 | 100 |
| NSP3_cd21732_betaCoV_PLPro | 100 | 100 | 100 | 100 |
| NSP3_cd21822_SARS-CoV-like_Nsp3_NAB | 99,9092 | 98,32 | 98,32 | 98 |
| NSP3_cl00019_Macro_SF | 99,9773 | 100 | 97,62 | 99 |
| NSP3_cl13138_SUD-M | 99,8865 | 99,17 | 96,77 | 98 |
| NSP3_cl13772_DUF3655 | 100 | 100 | 100 | 100 |
| NSP5_protein | 100 | 100 | 100 | 100 |
| NSP7_protein | 100 | 100 | 100 | 100 |
| NSP8_protein | 100 | 100 | 100 | 100 |
| NSP9_protein | 99,9319 | 98,22 | 100 | 99 |
| orf3a_protein | 99,9773 | 100 | 96,67 | 98 |
| orf7a_protein | 99,9546 | 100 | 92,31 | 96 |
| orf8_protein | 99,9773 | 95 | 100 | 97 |
| orf9b_protein | 100 | 100 | 100 | 100 |
| S_protein | 100 | 100 | 100 | 100 |
| **Average** | 99,9757 | 99,4157 | 99,1168 | 99,25 |

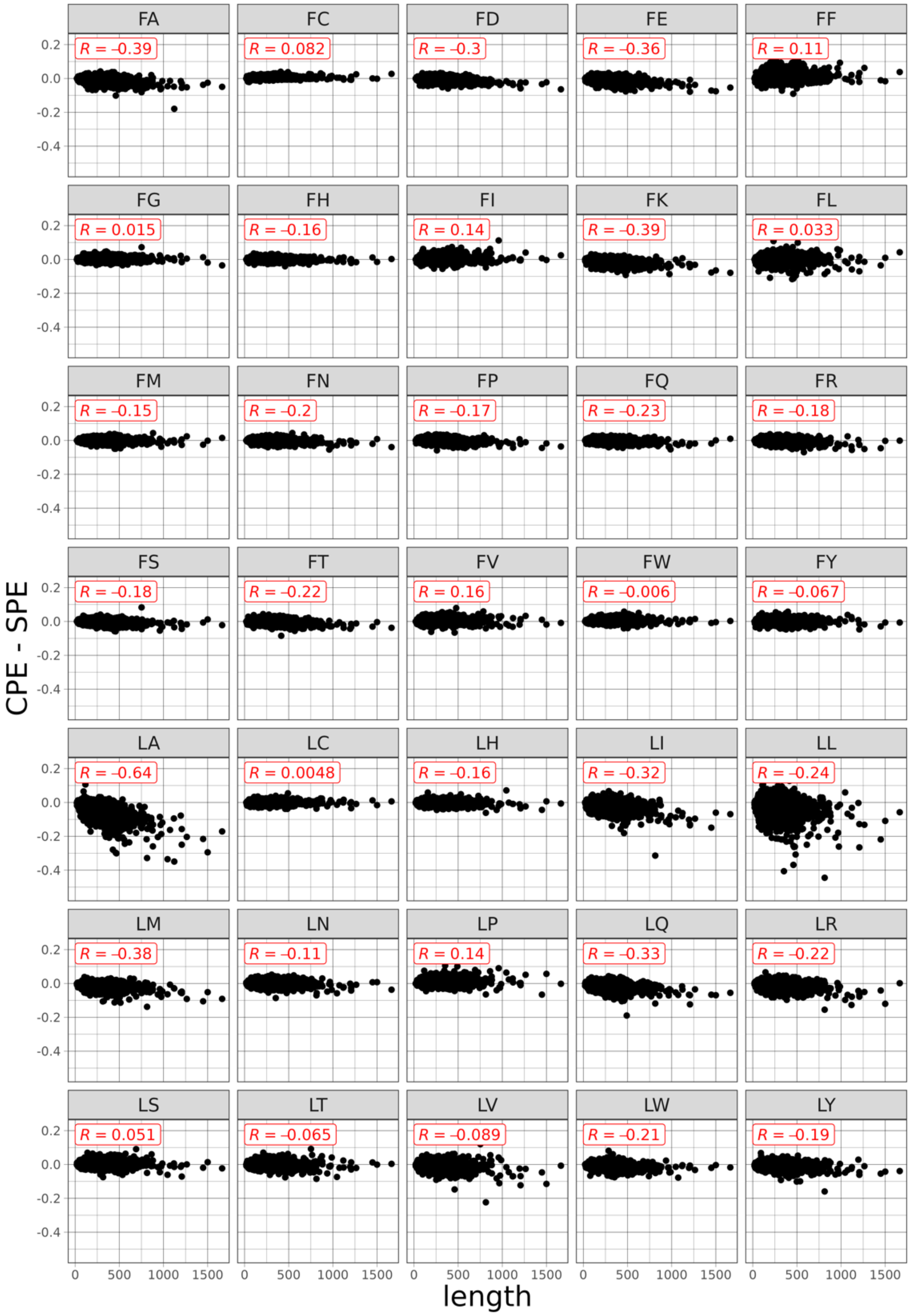


**Supplementary Fig. 1:** Scatter plots showing the differences in energy estimates (derived from sequence and structure) as a function of protein length across all 210 pairwise interactions, along with the correlation coefficients between these differences in energy estimates and protein length for each of the 210 pairwise interactions. Source data are provided as a Source Data file.

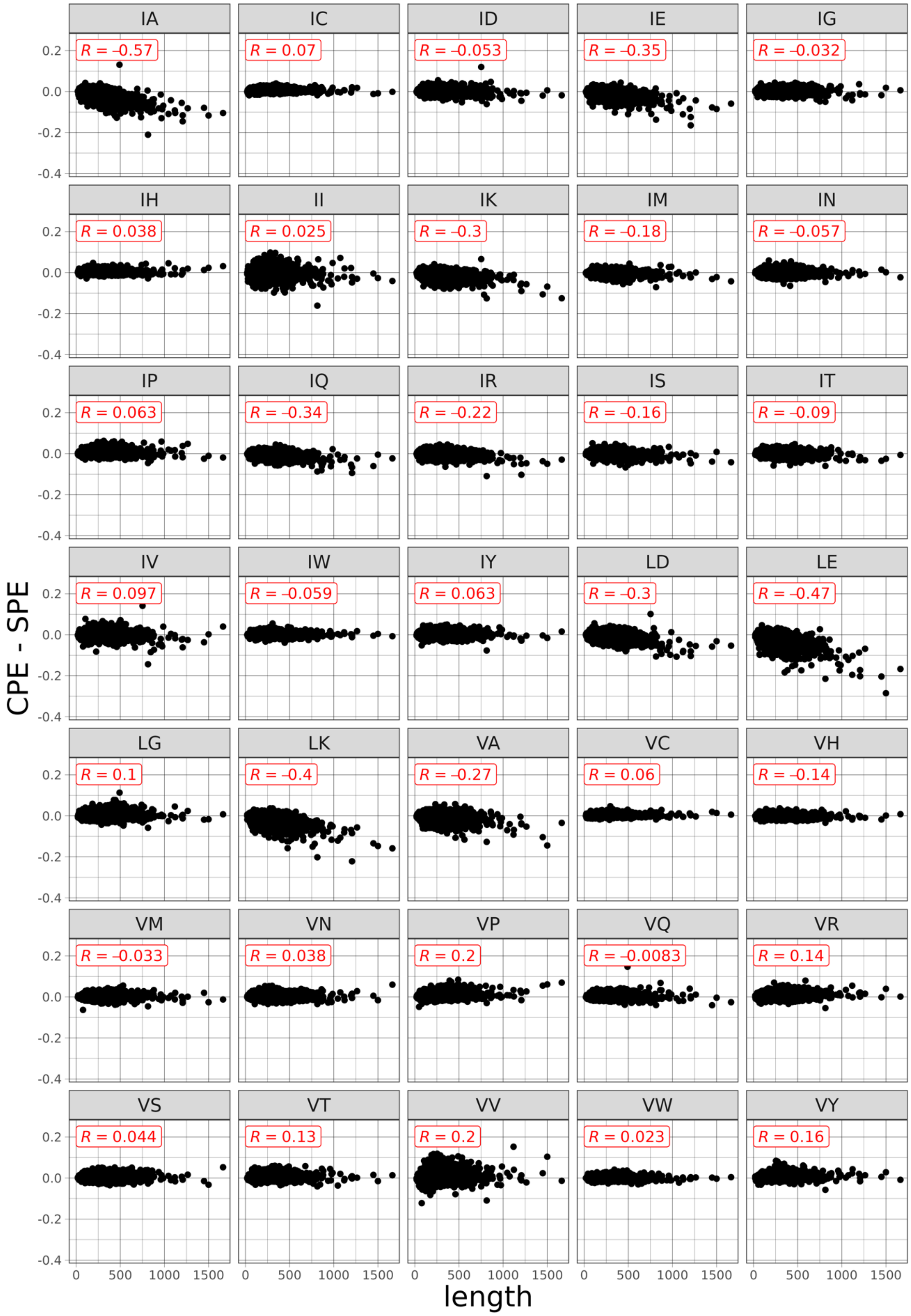


**Supplementary Fig. 1** (continued)

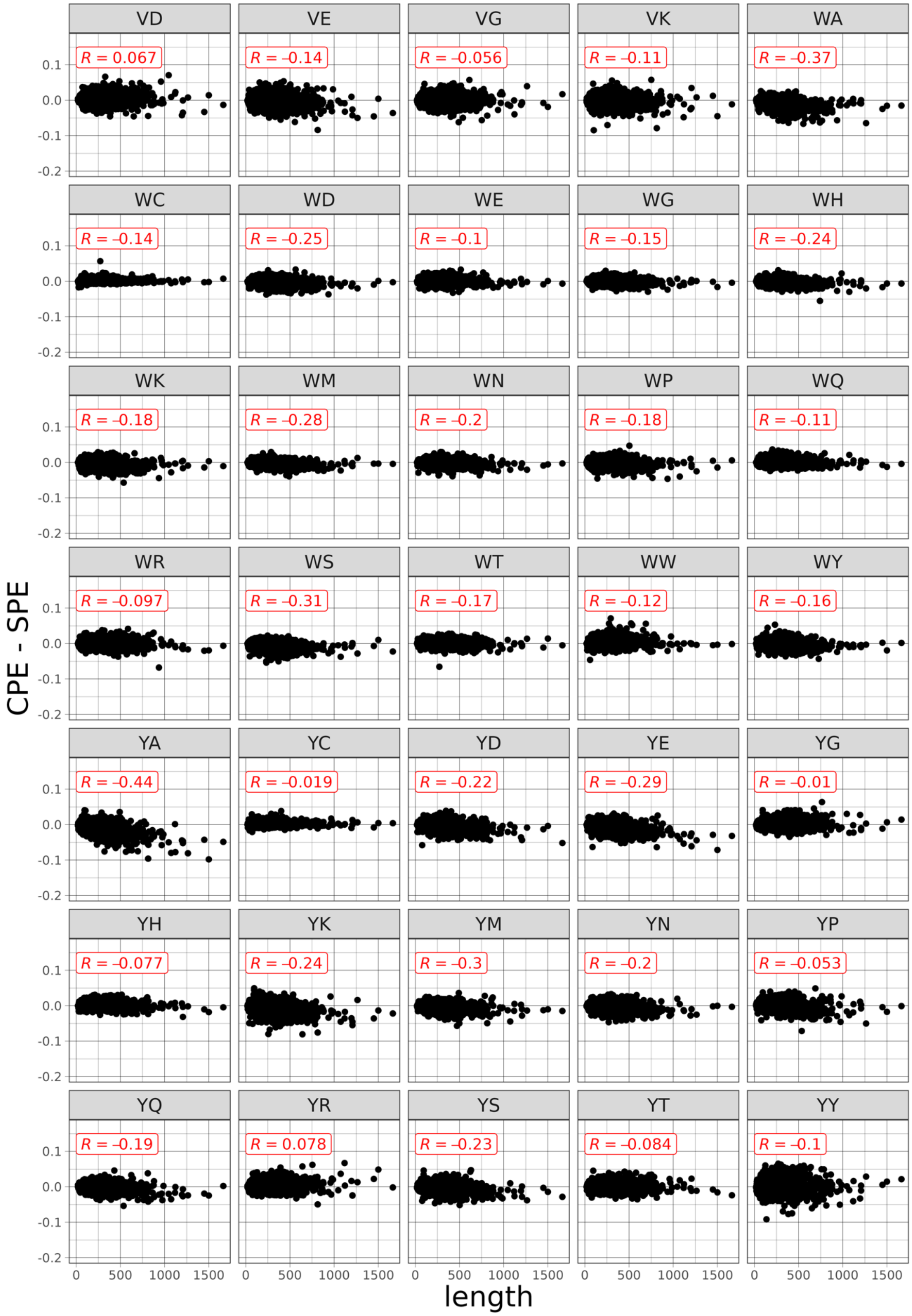


**Supplementary Fig. 1** (continued)

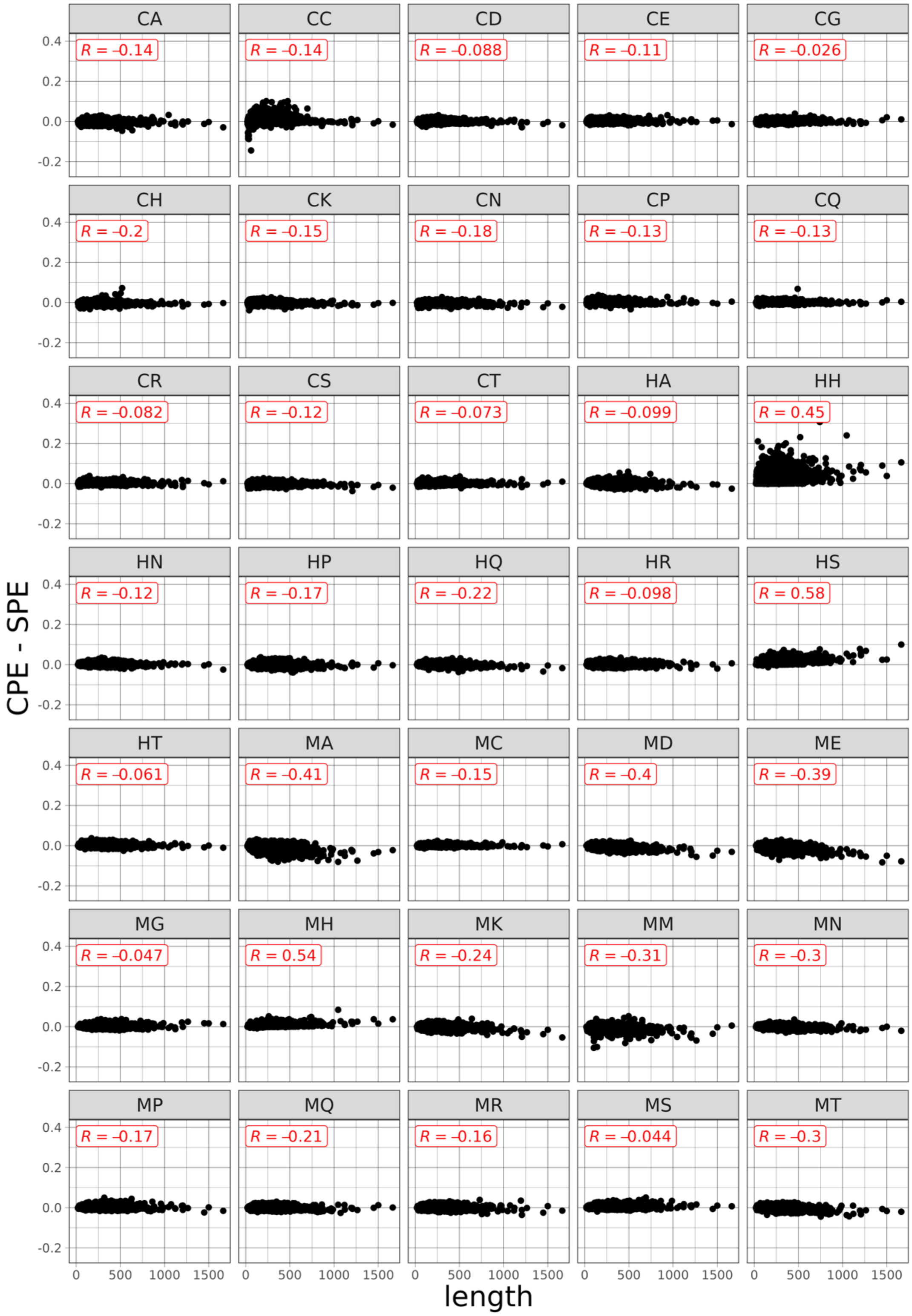


**Supplementary Fig. 1 (continued)**

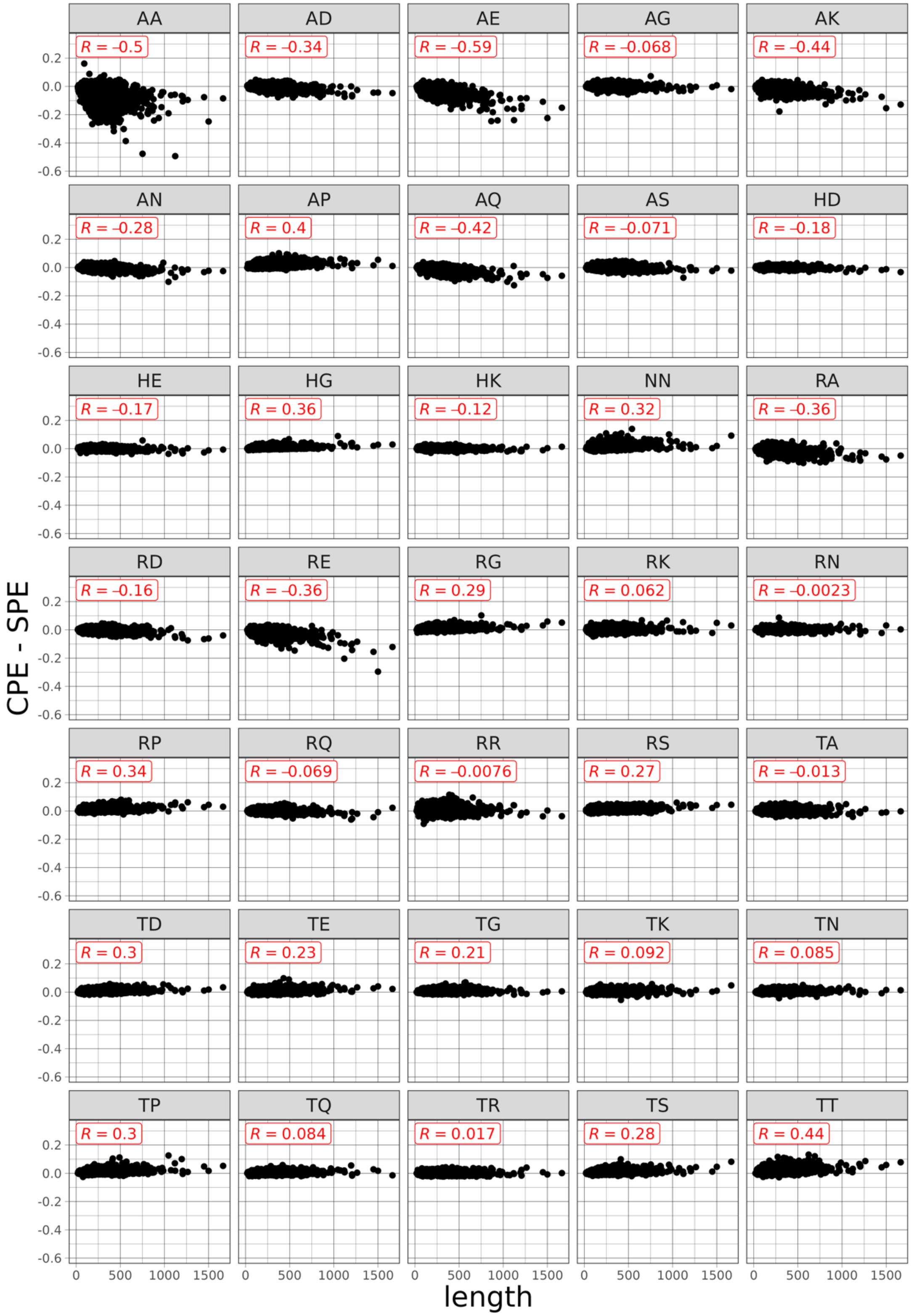


**Supplementary Fig. 1** (continued)

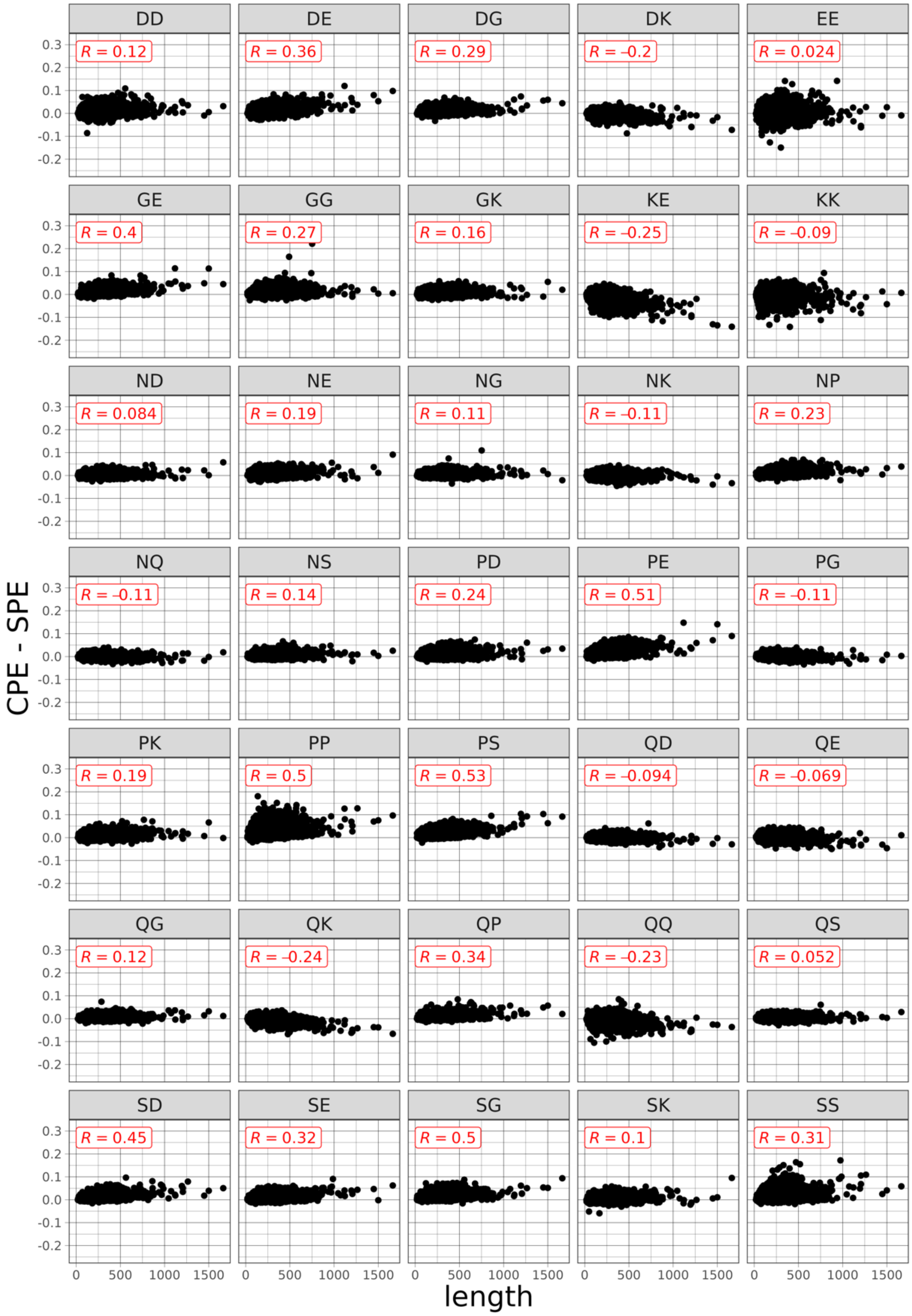


**Supplementary Fig. 1** (continued)

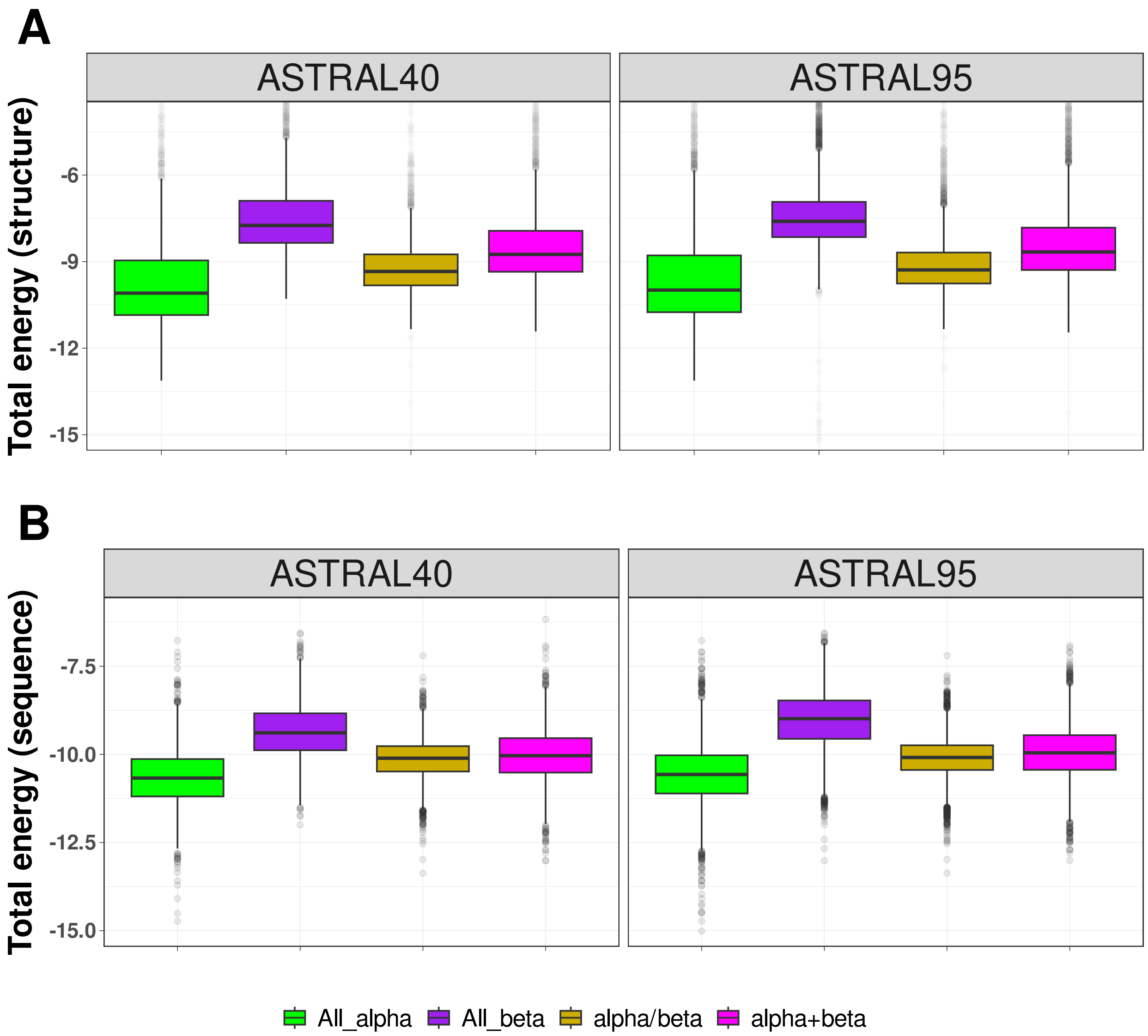


**Supplementary Fig. 2: Energy Distribution Across SCOP Structural Classes.** The distribution of normalized total energy in protein domains from ASTRAL40 and ASTRAL95 datasets based on protein structure (A) and sequence (B) across structural SCOP classes. In the ASTRAL40 dataset, there are 2643, 3058, 4462, and 3653 protein domains in the all-alpha, all-beta, alpha/beta, and alpha+beta classes, respectively. Similarly, in the ASTRAL95 dataset, there are 5442, 10160, 9342, and 7392 protein domains in the all-alpha, all-beta, alpha+beta, and alpha/beta classes, respectively. Boxplots display the median (center line), the 25th and 75th percentiles (bounds of the box), and the minimum and maximum values. Source data are provided as a Source Data file.

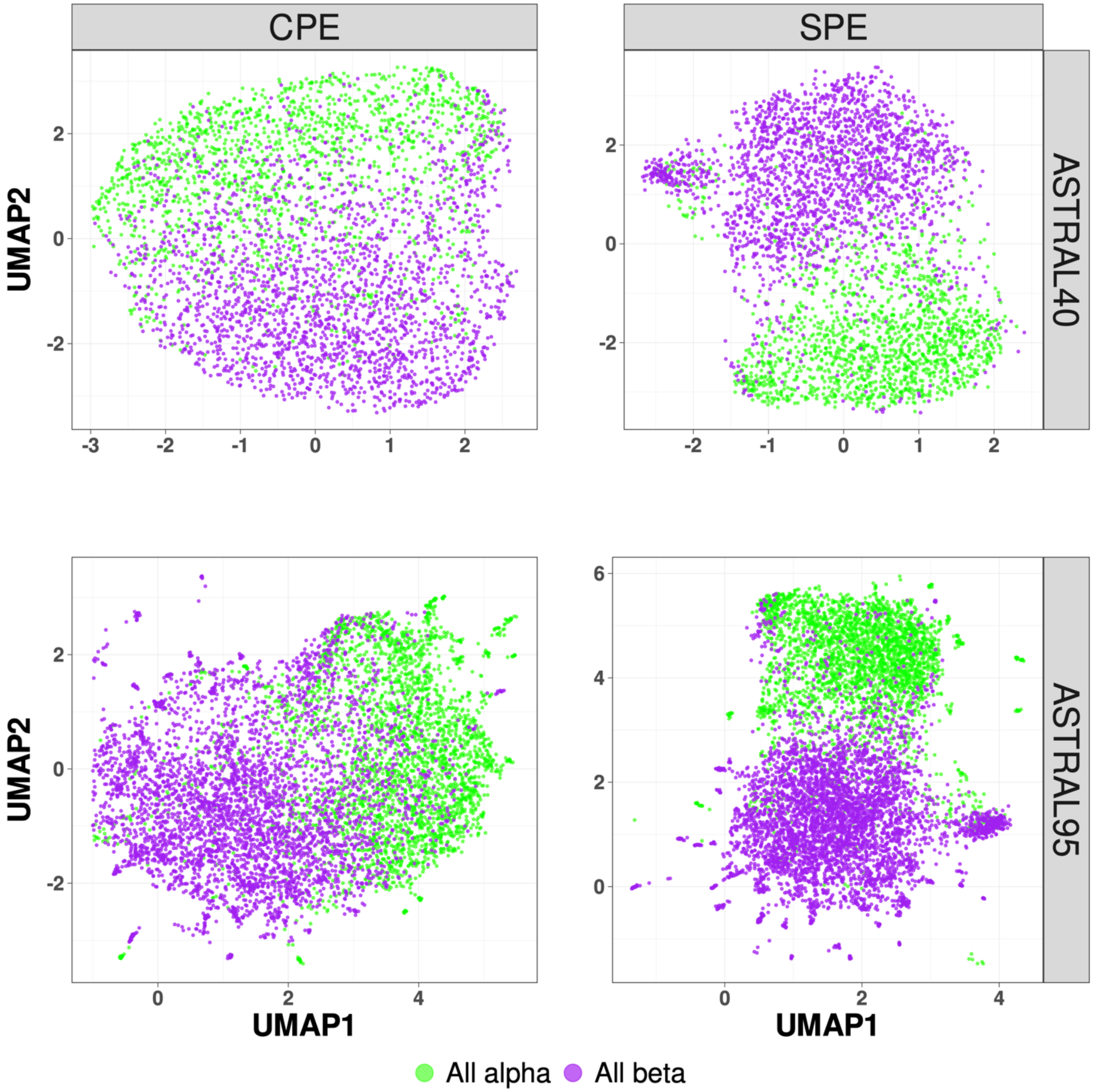


**Supplementary Fig. 3: UMAP Visualization of Energy Profiles in All-Alpha and All-Beta Domains from ASTRAL40 and ASTRAL95 Datasets.** UMAP projections of Sequence Profile Energy (SPE) and Contact Profile Energy (CPE) display the distribution of all-alpha (green) and all-beta (pink) proteins in the ASTRAL40 (top) and ASTRAL95 (bottom) datasets. Dots represent two dimensional UMAP projection of SPE(CPE) for individual sequences. UMAP plots were generated by parameters n_neighbors = 30 and min_dist = 0.1. Source data are provided as a Source Data file.

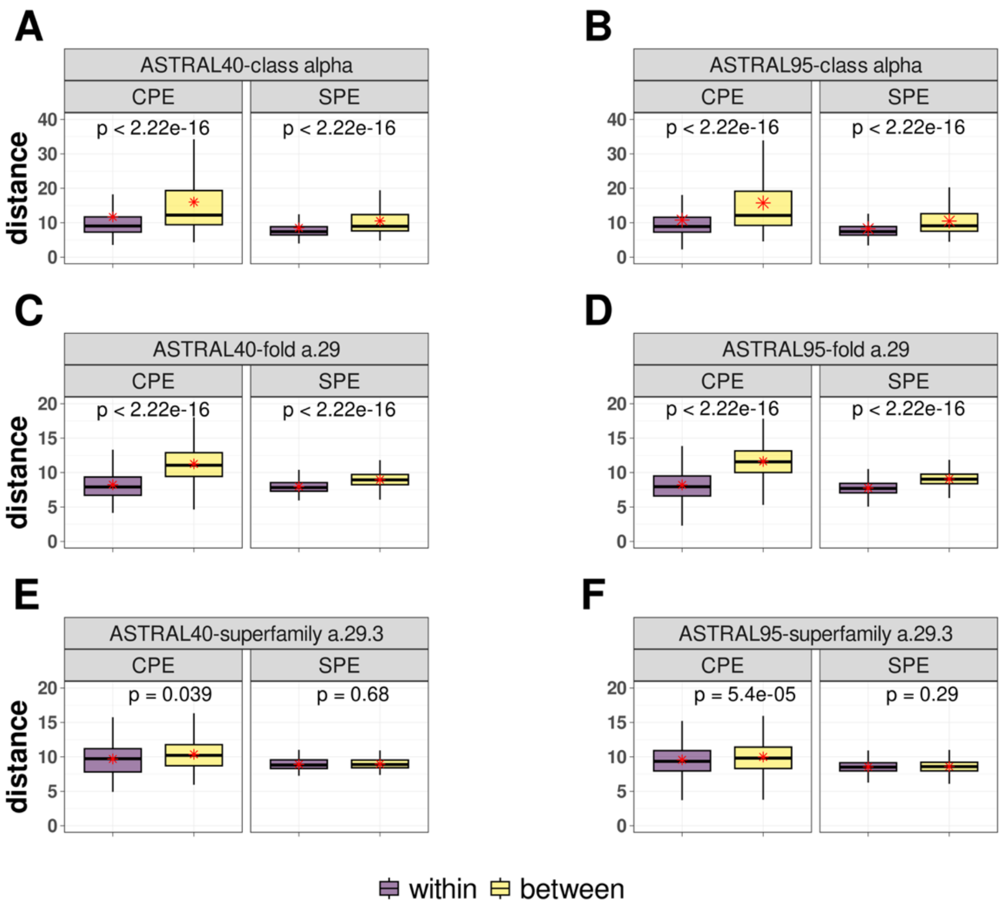


**Supplementary Fig. 4: Within-group and between-group distance comparisons using CPE and SPE methods across different SCOP levels in the ASTRAL dataset.** Boxplots depict the profile of energy distances for within-group (purple) and between-group (yellow) comparisons. A) Within-group distances represent pairwise comparisons among n= 1000 randomly selected pair domains from the class alpha, while between-group distances include comparisons between n=1000 alpha-class pair domains and domains from other classes in the ASTRAL40 dataset. B) The corresponding analysis with the same number of protein domains for the ASTRAL95 dataset. C) Within-group distances are derived from pairwise comparisons within the a.29 fold, comprising n= 650 pair domains, while between-group distances include comparisons with n= 3956 pair domains from different folds in the all-alpha class in ASTRAL40. D) Similarly, within-group comparisons involve n= 3872 pair domains in the a.29 fold, and between-group comparisons include n= 16748 pair domains from other folds in the all-alpha class in ASTRAL95. E) Within-group comparisons are within the a.29.3 superfamily (77 protein domains), while between-group comparisons include n= 188 pair domains from other superfamilies within the a.29 fold in ASTRAL40. F) Within-group comparisons in the a.29.3 superfamily involve n= 813 pair domains, and between-group comparisons involve n= 1344 pair domains from other superfamilies within the a.29 fold in ASTRAL95. Statistics assessed by two-tailed Student's t-test. P-values (indicated above the boxplots) assess the significance of the differences between within- and between-group distances. Boxplots display the median (center line), the 25th and 75th percentiles (bounds of the box), and the minimum and maximum values (whiskers) excluding outliers. The exact p-value is less than 10e-16, which is below the precision threshold of standard statistical computations. Source data are provided as a Source Data file.

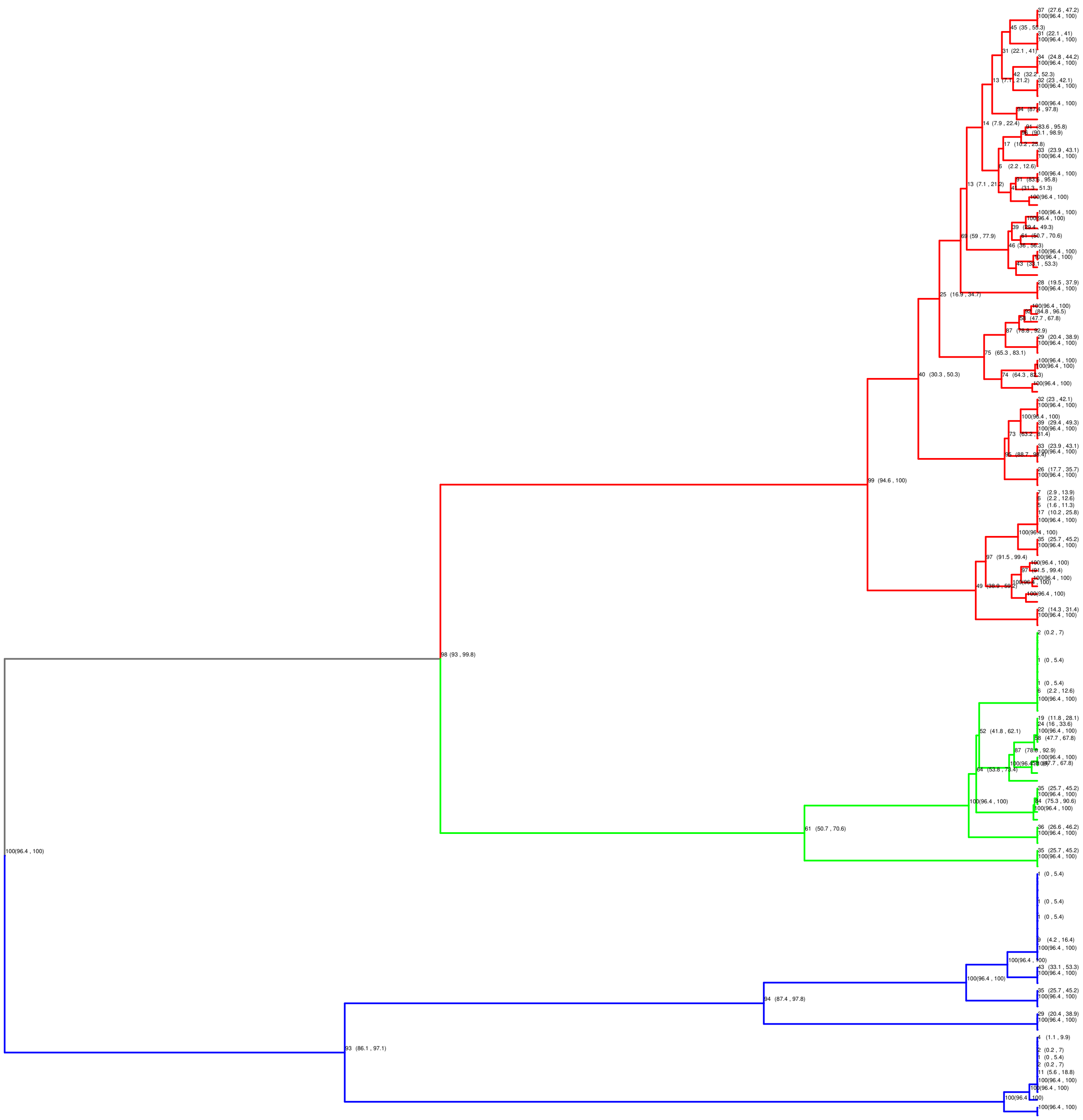

**Supplementary Fig. 5:** Bootstrap and confidence interval analysis for phylogenetic tree reconstruction using spike glycoprotein structures of SARS-CoV, SARS-CoV-2, and MERS-CoV based on the CPE method.

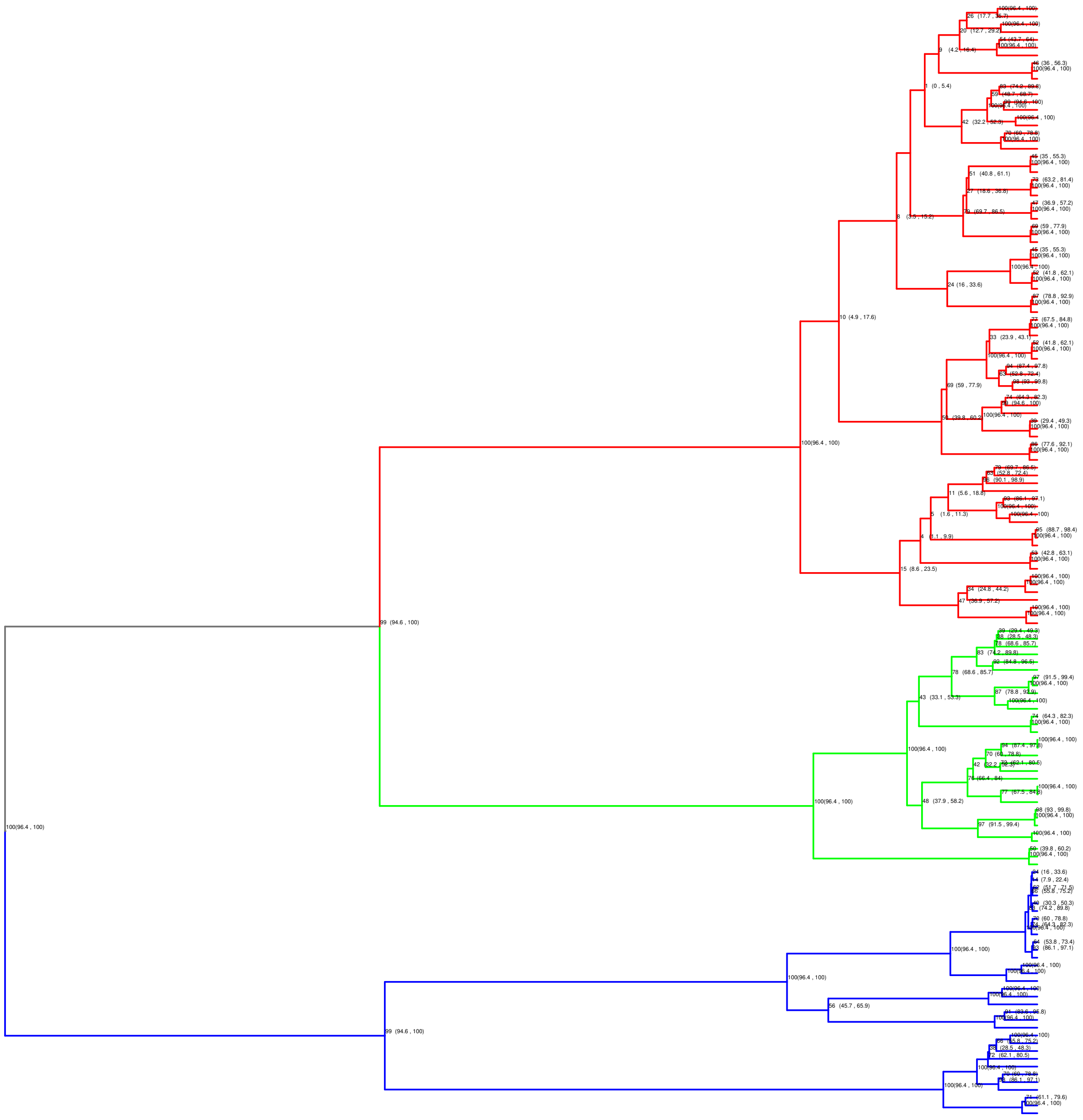

**Supplementary Fig. 6:** Bootstrap and confidence interval analysis for phylogenetic tree reconstruction using spike glycoprotein structures of SARS-CoV, SARS-CoV-2, and MERS-CoV based on the SPE method.

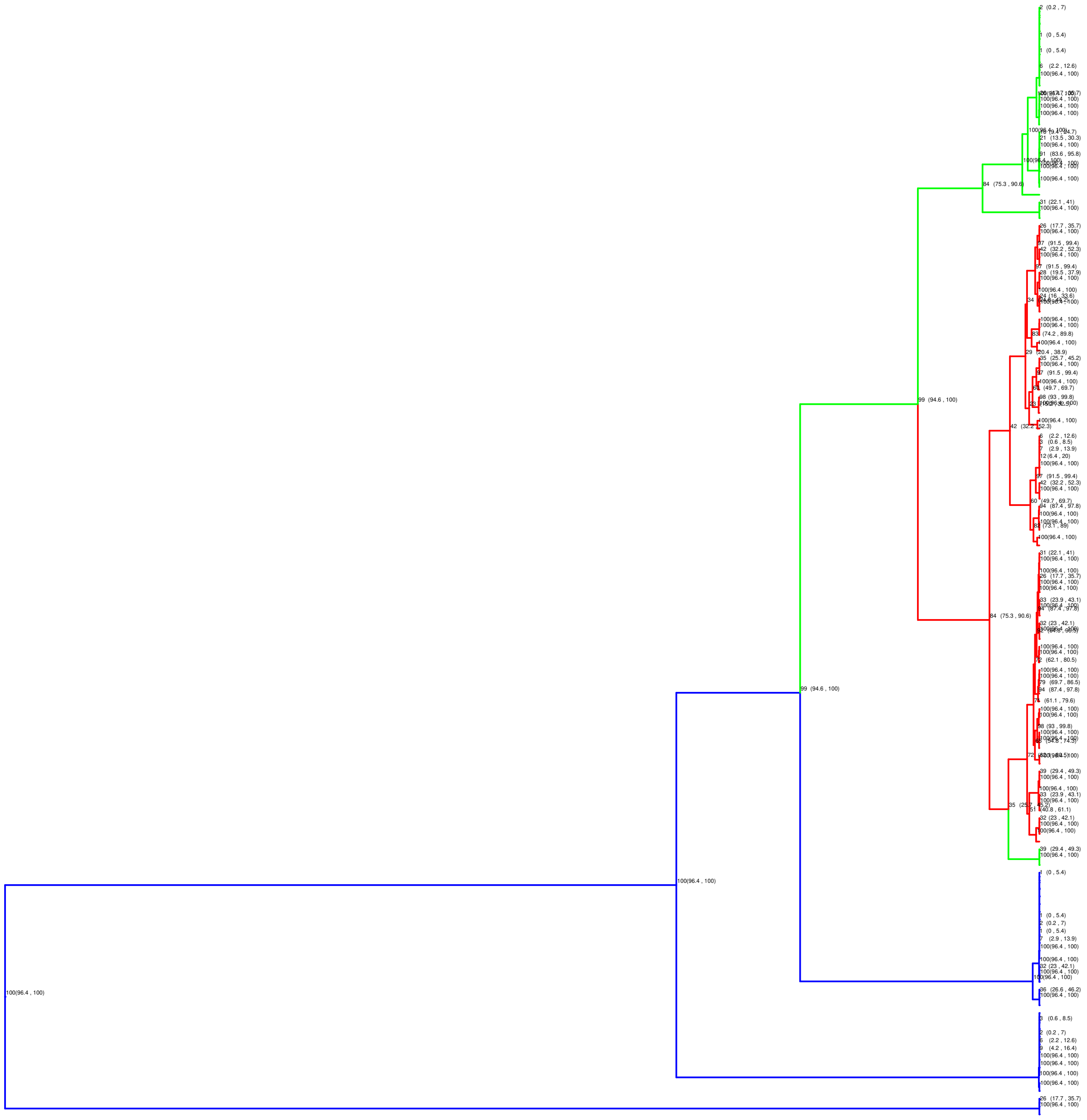

**Supplementary Fig. 7:** Bootstrap and confidence interval analysis for phylogenetic tree reconstruction using spike glycoprotein structures of SARS-CoV, SARS-CoV-2, and MERS-CoV based on the TM-Vec method.

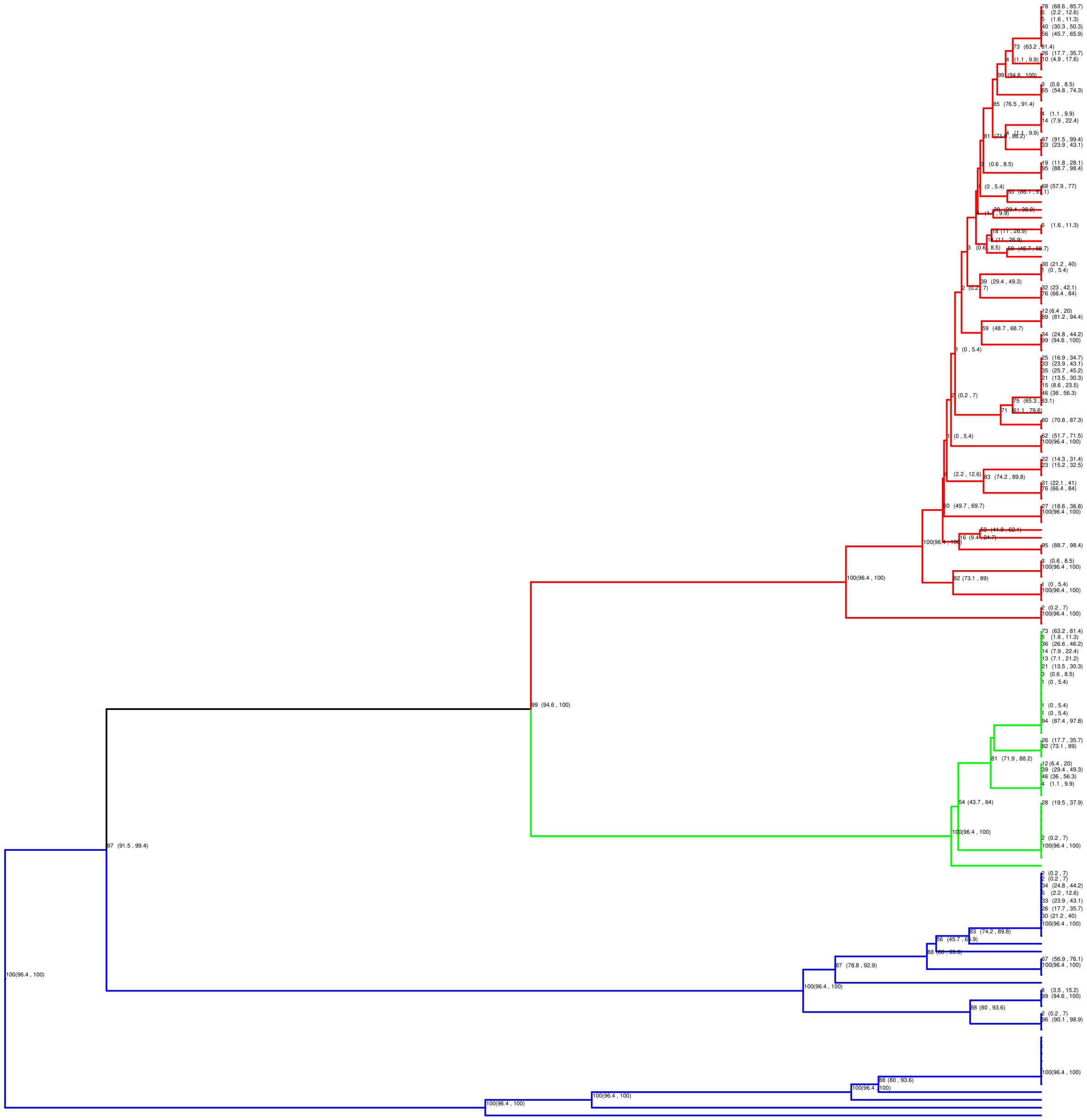

**Supplementary Fig. 8:** Bootstrap and confidence interval analysis for phylogenetic tree reconstruction using spike glycoprotein structures of SARS-CoV, SARS-CoV-2, and MERS-CoV based on the MSA method.

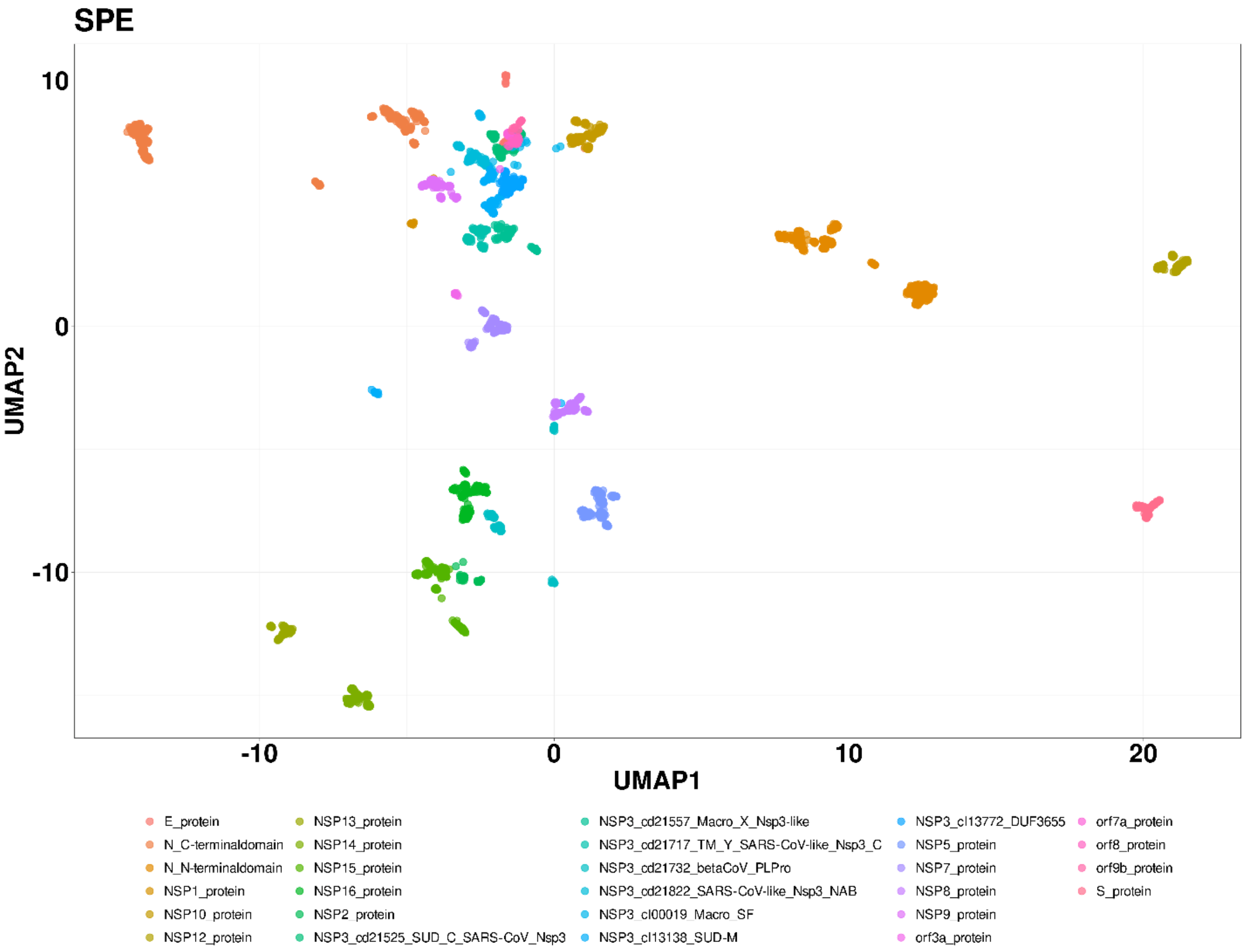


**Supplementary Fig. 9: UMAP Visualization of Energy Profiles in Large-Scale SARS-Cov2 data set.** The UMAP projection of Structural Energy Profiles (SPE) on 28 protein families with a total of 4,405 protein models. UMAP plot was generated using parameters n_neighbors = 150 and min_dist = 0.1. Source data are provided as a Source Data file.

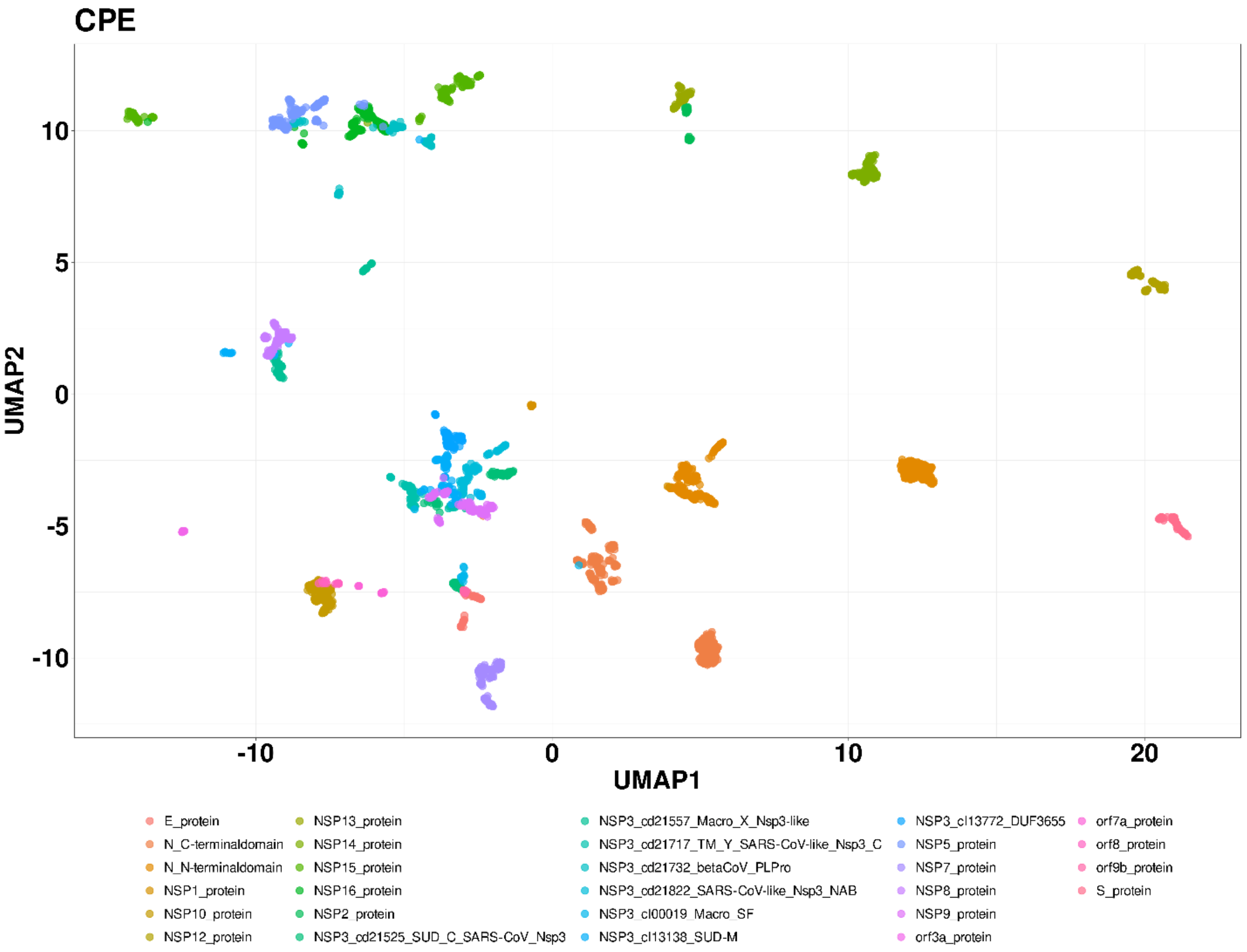


**Supplementary Fig. 10: UMAP Visualization of Energy Profiles in Large-Scale SARS-Cov2 data set.** Th e UMAP projection of Compositional Energy Profiles (CPE) on 28 protein families with a total of 4,405 protein models. UMAP plot was generated using parameters n_neighbors = 150 and min_dist = 0.1. Source data are provided as a Source Data file.

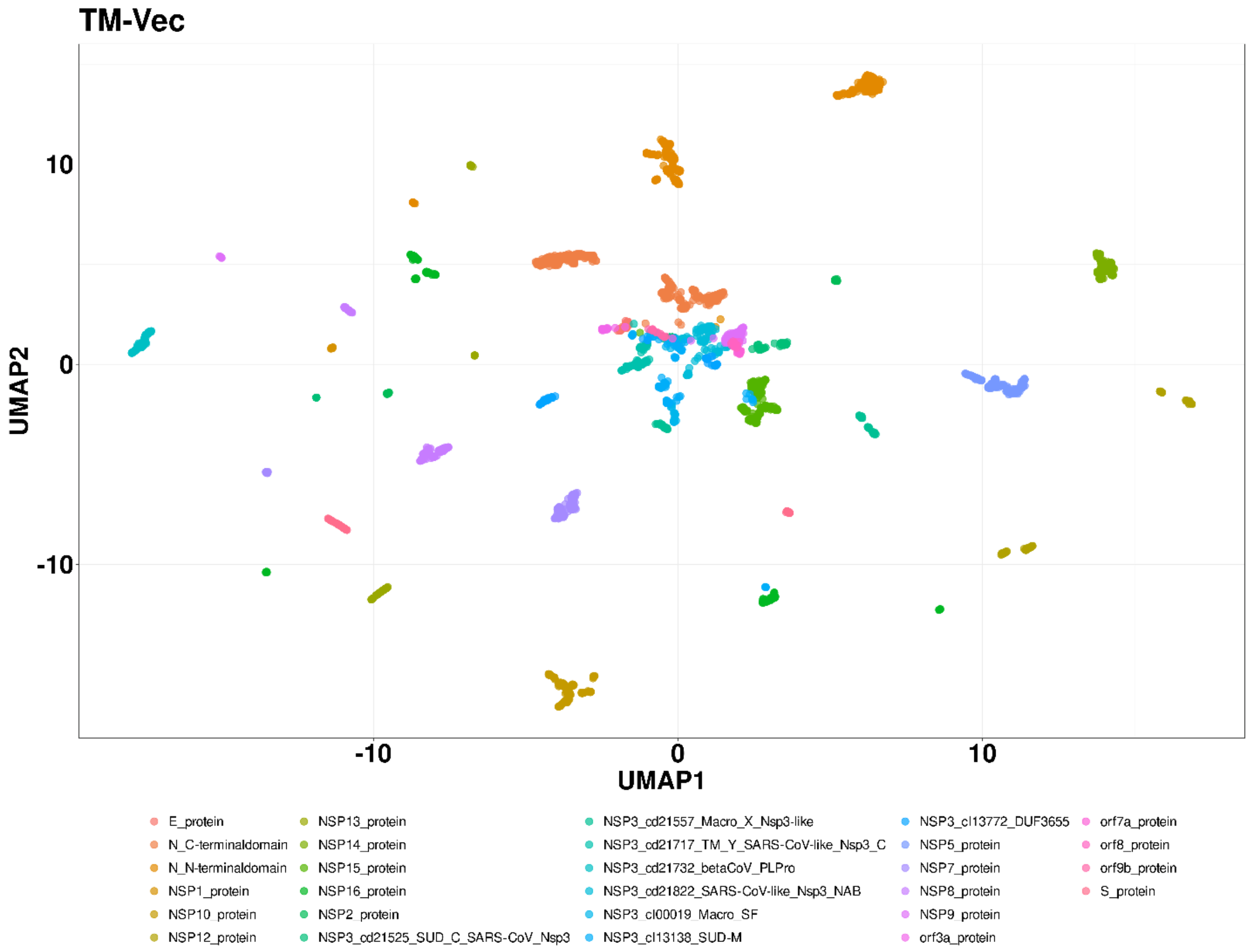


**Supplementary Fig. 11: UMAP Visualization of Energy Profiles in Large-Scale SARS-Cov2 data set.** The UMAP projection of TM-Vec on 28 protein families with a total of 4,405 protein models. UMAP plot was generated using parameters n_neighbors = 150 and min_dist = 0.1. Source data are provided as a Source Data file.
